# Disparate temperature-dependent virus – host dynamics for SARS-CoV-2 and SARS-CoV in the human respiratory epithelium

**DOI:** 10.1101/2020.04.27.062315

**Authors:** Philip V’kovski, Mitra Gultom, Jenna Kelly, Silvio Steiner, Julie Russeil, Bastien Mangeat, Elisa Cora, Joern Pezoldt, Melle Holwerda, Annika Kratzel, Laura Laloli, Manon Wider, Jasmine Portmann, Thao Tran, Nadine Ebert, Hanspeter Stalder, Rune Hartmann, Vincent Gardeux, Daniel Alpern, Bart Deplancke, Volker Thiel, Ronald Dijkman

## Abstract

Since its emergence in December 2019, SARS-CoV-2 has spread globally and become a major public health burden. Despite its close phylogenetic relationship to SARS-CoV, SARS-CoV-2 exhibits increased human-to-human transmission dynamics, likely due to efficient early replication in the upper respiratory epithelium of infected individuals. Since different temperatures encountered in the human respiratory tract have been shown to affect the replication kinetics of several viruses, as well as host immune response dynamics, we investigated the impact of temperatures during SARS-CoV-2 and SARS-CoV infection in the human airway epithelial cell culture model. SARS-CoV-2, in contrast to SARS-CoV, replicated more efficiently at temperatures encountered in the upper respiratory tract, and displayed higher sensitivity to type I and type III IFNs. Time-resolved transcriptome analysis highlighted a temperature-dependent and virus-specific induction of the IFN-mediated antiviral response. These data reflect clinical features of SARS-CoV-2 and SARS-CoV, as well as their associated transmission efficiencies, and provide crucial insight on pivotal virus - host interaction dynamics.

## Introduction

In December 2019, a new zoonotic coronavirus emerged in Wuhan, Hubei Province, China, which is referred to as Severe Acute Respiratory Syndrome Coronavirus 2 (SARS-CoV-2), and is the etiological agent of Coronavirus Disease 2019 (COVID-19) ^1–3^. The novel coronavirus has a close phylogenetic relationship with SARS-CoV, which emerged in China in 2002/2003 and led to over 8000 confirmed cases worldwide, including 800 deaths ^4^. SARS-CoV-2 differs from SARS-CoV by only 380 amino acids over its entire 30 kb genome and retains a high level of conservation in the receptor binding motifs that interact with the human receptor angiotensin converting enzyme 2 (ACE2) ^5^. Despite these similarities, there are currently over 40 million confirmed cases of SARS-CoV-2 worldwide, including more than 1 million deaths ^6^. Moreover, although the cell surface receptor ACE2 and the serine protease TMPRSS2 have been demonstrated to serve as entry determinants for both SARS-CoV and SARS-CoV-2 ^2,7–9^, an accumulating body of evidence showing dissimilar human-to-human transmission dynamics and clinical courses between SARS-CoV-2 and SARS-CoV strongly suggests the presence of disparate virus-host dynamics during viral infection in the human respiratory epithelium ^10–15^.

The human conductive respiratory tract is lined by a pseudostratified, ciliated, columnar epithelium that contains mucin-producing goblet cells and represents a crucial barrier to constrain invading pathogens. The anatomical distance between the upper and lower respiratory conductive tract, and their different ambient temperatures (32-33°C and 37°C, respectively ^16,17^), have previously been shown to influence the replication kinetics of diverse respiratory viruses, such as rhinoviruses, influenza viruses and coronaviruses ^18–22^. Moreover, the anatomical disparity in ambient temperature also affects virus – host immune response dynamics, and thus potential human-to-human transmission dynamics ^23^. Interestingly, SARS-CoV-2 has been detected earlier after infection than SARS-CoV in upper respiratory tissues of infected patients ^10,13,14,24,25^, suggesting that transmission kinetics and host innate immune response dynamics might differ between SARS-CoV and SARS-CoV-2 infections.

Since viral load may reflect the dynamic interaction between viral replication and inhibition by cellular defence mechanisms, we employed the human airway epithelial cell (hAEC) culture model to investigate the influence of different incubation temperatures on the viral replication kinetics and host immune response dynamics of both SARS-CoV and SARS-CoV-2 infections. Our study revealed that SARS-CoV-2 replication improved in hAEC cultures incubated at 33°C rather than 37°C and resulted in higher titers than SARS-CoV, while both viruses replicated equally efficiently at 37°C. Pretreatment of hAEC cultures with exogenous type I and III interferon (IFN) at different temperatures showed that SARS-CoV-2 and SARS-CoV are equally sensitive to both type I and III IFN, thereby exemplifying the relevance of early IFN signalling and innate immune responses to restrict viral infection. Importantly, a detailed temporal transcriptome analysis of infected hAEC cultures corroborated initial findings and uncovered characteristic innate immune response gene signatures relating to the viral replication efficiency of SARS-CoV and SARS-CoV-2 at different ambient temperatures. Altogether, these results provide the first in-depth fundamental insight on the virus-host innate immune response dynamics of SARS-CoV-2 and the closely phylogenetically related SARS-CoV in the respiratory epithelium, and likely reflect the clinical characteristics and transmission efficiencies of both viruses.

## Results

### Replication kinetics of SARS-CoV-2 and SARS-CoV at 33°C and 37°C

To assess the influence of the temperature variations that occur along the human respiratory tract, and to model virus-host interaction dynamics in distinct anatomical regions, we maintained well-differentiated hAEC cultures at either 33°C or 37°C throughout the experiment. hAEC cultures represent a well-characterized *in vitro* model that morphologically and functionally recapitulate the epithelial lining of the human respiratory tract *in vivo*. hAEC cultures from seven different human donors were inoculated with either SARS-CoV-2/München-1.1/2020/929 or SARS-CoV Frankfurt 1 isolates using a multiplicity of infection (MOI) of 0.1. The polarity of viral progeny release was monitored by collecting apical washes and basolateral medium in 24-hour intervals for a period of 96 hours. Since SARS-CoV-2 can be detected early after infection in the upper respiratory specimens of infected patients ^14,24,25^, we incubated virus-infected cultures at both 33°C and 37°C to mimic the ambient temperatures of the human upper and lower respiratory tract, respectively. At 37°C SARS-CoV and SARS-CoV-2 replicated to similar titers over the course of the infection (**Fig. 1a, b**). Interestingly, when assessing viral replication efficiency at 33°C rather than at 37°C, it was apparent that SARS-CoV-2 infection resulted in 10-fold higher titers released in the apical compartment between 72 and 96 hours post infection (hpi). In contrast, SARS-CoV replication at 33°C remained similar to replication at 37°C and showed no significant differences over the entire course of infection (**Fig. 1a, b**). Since the directionality of viral progeny release is crucial for subsequent virus spread and overall disease outcome, we also assessed whether SARS-CoV-2 was released to the apical surface, basolateral surface, or bilaterally. Similar to what we and others have observed previously for all other human coronaviruses, SARS-CoV-2 was predominantly released to the luminal surface (**Supp. Fig. 1a, b**) ^26,27^.

**Figure 1.**
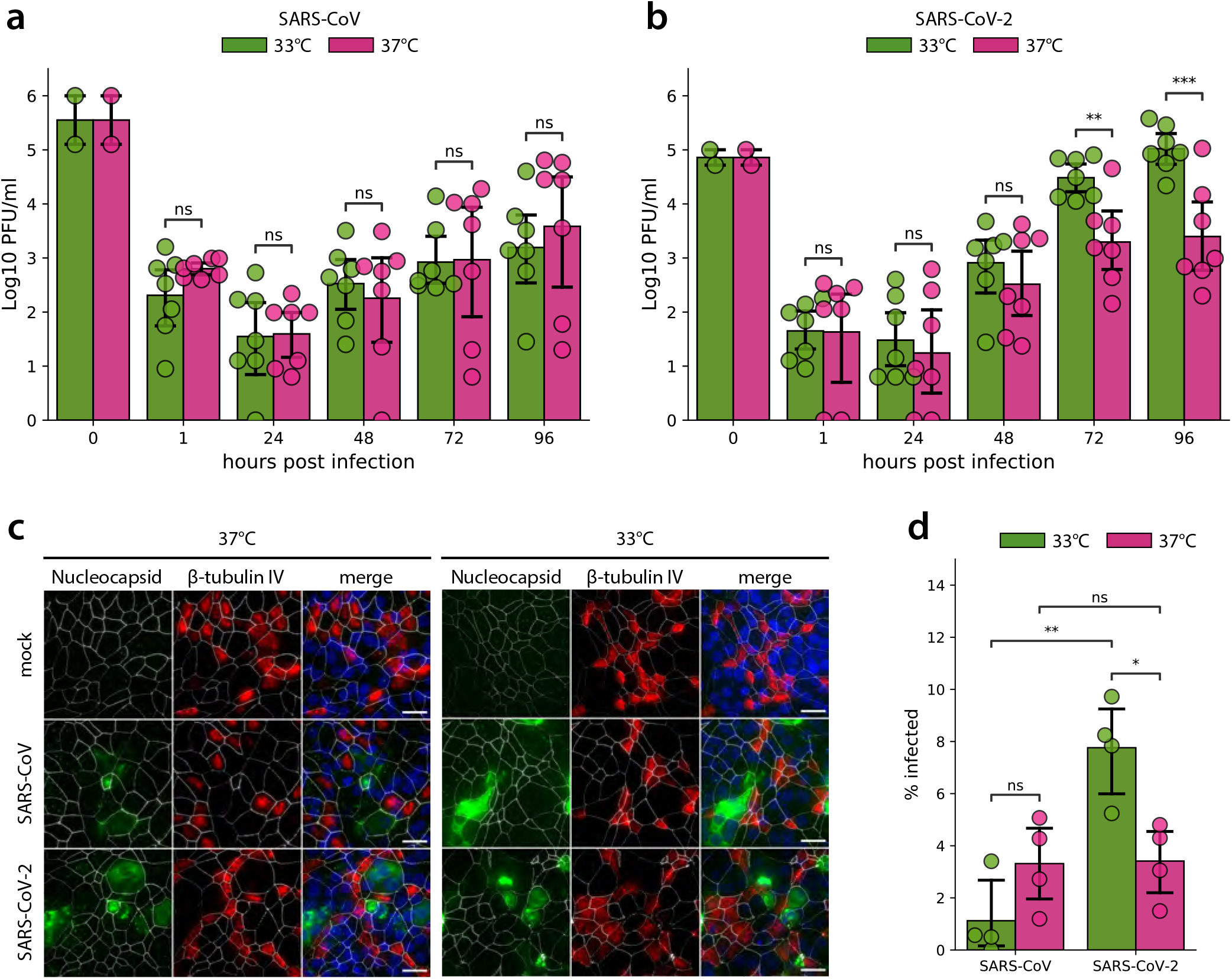
SARS-CoV and SARS-CoV-2 replication kinetics in hAEC cultures. Well-differentiated hAEC cultures were infected with SARS-CoV (**a**) and SARS-CoV-2 (**b**) using 30,000 PFU or remain uninfected (mock), and were incubated at 37°C or 33°C. Inoculated virus was removed at 1 hpi and the apical side was washed. Cultures were further incubated at the indicated temperature. At the indicated time post infection, apical virus release was assessed by plaque titration (**a-b**). Data represent the mean ± 95% CI of hAEC cultures from seven different human donors. Individual points represent the average of two technical replicates. Values at 0 hpi indicate the titer of the inoculum used to infect the hAEC cultures, and values at 1 hpi indicate the remaining titer after the third wash. The p-values were computed by using two-sided paired sample t tests. At 96 hpi, hAEC cultures were fixed and processed for immunofluorescence analysis using antibodies against SARS-CoV Nucleocapsid protein (green), β-tubulin (cilia, red), ZO-1 (tight junctions, white) and DAPI (blue) (**c**). Representative z-projections of one donor are shown. Scale bar, 20 microns. Infected cells were quantified after segmentation of individual cells based on the ZO-1 staining and measuring the mean intensity of the nucleocapsid protein staining (**d**). Data represent the mean ± 95% CI of multiple images acquired from hAEC cultures derived from four different human donors. On average, more than 10^4^ cells per donor and per condition were analysed.

To assess whether the observed differential temperature-dependent replication efficiencies are a result of the number of cells infected by SARS-CoV or SARS-CoV-2, hAEC cultures were fixed at 96 hpi and processed for immunofluorescence analysis using antibodies directed against the SARS-CoV Nucleocapsid protein. Additionally, to discern potential preferential virus tropism to a distinct cell type, characteristic markers of the hAEC culture’s architecture, such as the intercellular tight junctions (ZO-1) and the presence of cilia (β-tubulin), were also included. Microscopy investigations and automated image quantification revealed that in accordance with the more efficient replication kinetics of SARS-CoV-2 at 33°C, the fraction of SARS-CoV-2 infected cells increased significantly at 33°C compared to 37°C and to SARS-CoV. On the other hand, SARS-CoV did not show any significant change in the fraction of infected cells at 33°C compared to 37°C. Both viruses display a comparable fraction of infected cells at 37°C. (**Fig. 1c, d**). Notably, at 96 hpi the majority of SARS-CoV and SARS-CoV-2 antigen positive cells were not co-stained by the β-tubulin marker and were therefore qualified as non-ciliated cells (**Fig. 1c**). We previously observed that SARS-CoV infects both non-ciliated and ciliated cell populations ^26^, however, given that other reports show that SARS-CoV primarily targets ciliated cells ^28^, we analysed the localization of the entry receptor, ACE2, and β-tubulin markers by microscopy. Immunofluorescence analysis revealed that ACE2 is expressed in both ciliated and non-ciliated cell populations in uninfected hAEC cultures (**Supp. Fig. 2a**). In line with this, analysis of mRNA expression in non-infected hAEC cultures using single-cell RNA-sequencing (scRNA-seq) confirmed that both *ACE2* and *TMPRSS2* mRNA are found in both secretory and ciliated cell populations (**Supp. Fig. 2b-d**) ^29^ Combined these results demonstrate that despite their shared requirement on ACE2 and TMPRSS2 for entry into host cells, SARS-CoV and SARS-CoV-2 display strong temperature-dependent variation in replication kinetics in hAEC cultures, suggestive of host determinants intervening during post-entry stages of the viral life cycle. Importantly, the significantly enhanced replication of SARS-CoV-2 at 33°C likely supports the increased replication in the upper respiratory tract and transmissibility of SARS-CoV-2 compared to SARS-CoV.

### Sensitivity of SARS-CoV-2 and SARS-CoV to IFN

The amount of viral progeny secreted from infected cells may reflect the dynamic interplay between viral replication and its inhibition by cellular defence mechanisms, such as by different types of interferon stimulated genes (ISGs). To examine whether the induction of ISGs differentially affects SARS-CoV and SARS-CoV-2 replication, hAEC cultures were pretreated with 50 IU or 5 IU of exogenous type I interferon (IFN-αA/D) and 50 or 5 ng of type III interferon (IFN-□3), for 18 hours prior to infection, at either 33°C or 37°C. Hereafter, the hAEC culture medium was replaced with IFN-free medium and cells were infected with SARS-CoV and SARS-CoV-2 at an MOI of 0.1, at 33°C or 37°C, for 72 hours. The titration of apically released virus revealed that the replication of both SARS-CoV-2 and SARS-CoV is severely restricted upon pretreatment with 50 IU of type I or 50 ng of III IFN at either 37°C or 33°C (**Fig. 2a**). Although, similar to previously reported observations ^26^, SARS-CoV seemed less sensitive to type I IFN than to type III IFN pretreatment at 37°C (**Fig. 2a, 24 hpi**), these differences were not statistically significant, which was further confirmed by pretreatment with 5 IU of type I IFN or 5 ng of type III IFN (**Fig. 2b**). Consistently, SARS-CoV displayed a similar sensitivity to type I and III IFN pretreatments at 33°C (**Fig. 2a, b**). In line with these results, SARS-CoV-2 was equally sensitive to both doses of type I or III IFN pretreatements at either 37°C or 33°C (**Fig. 2a, b**). However, overall SARS-CoV-2 replication kinetics were more severely impaired by IFN pretreatement compared to SARS-CoV. In contrast, restriction of SARS-CoV-2 upon pretreatement with 10 IU of type I and III IFNs was moderately relieved at 72 hpi (**Fig. 2b**).

**Figure 2.**
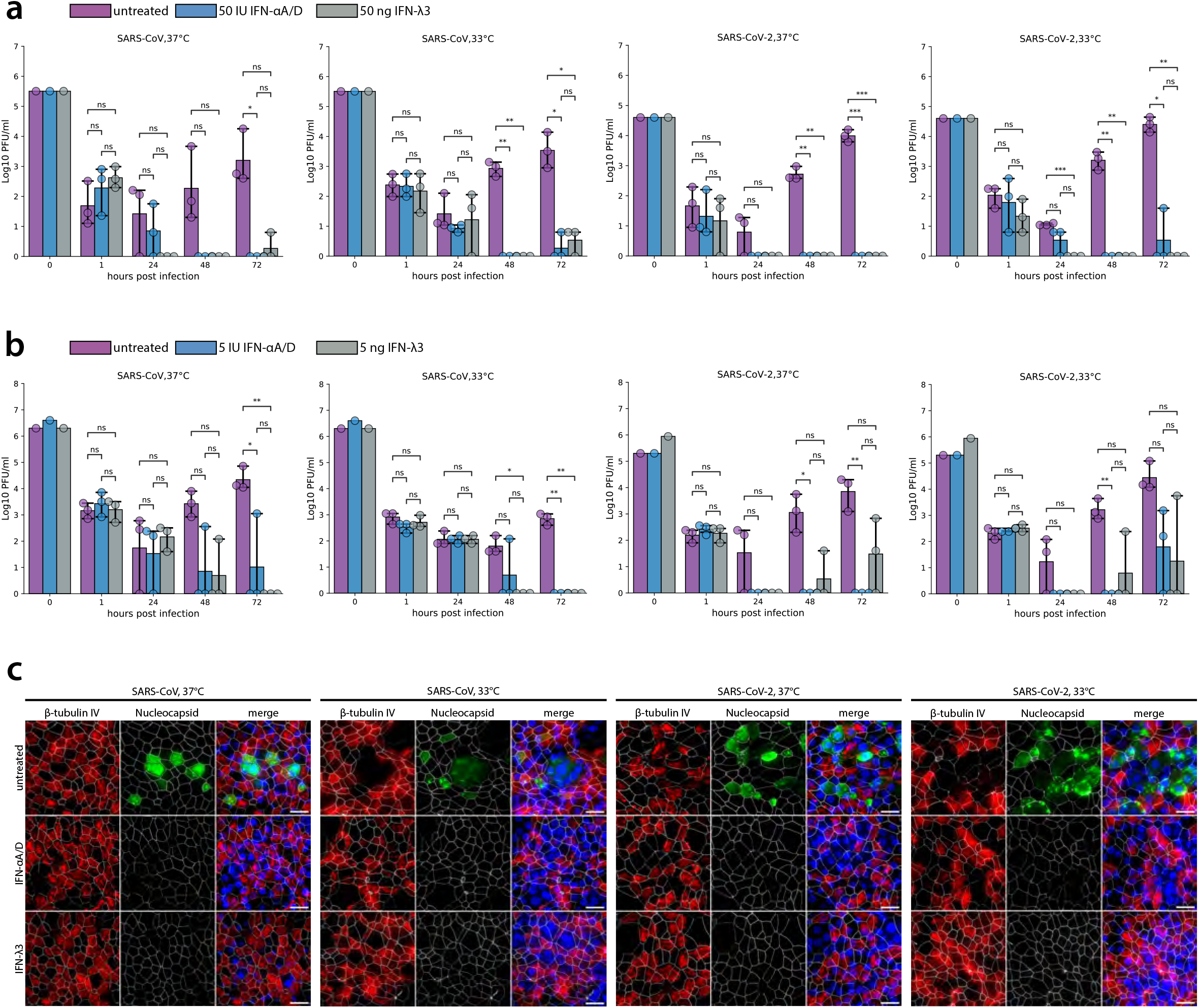
SARS-CoV and SARS-CoV replication upon IFN-I and IFN-III pretreatment. hAEC cultures were treated from the basolateral side with recombinant universal type I interferon (100 IU/ml or 10 IU/ml) or recombinant IFN-λ3 (100 ng/ml or 10 ng/ml) for 18 h. Before infection, medium was removed and replaced with IFN-free medium and hAEC cultures were infected with SARS-CoV and SARS-CoV-2 using 30,000 PFU, and were incubated at 37°C or 33°C. Inoculated virus was removed at 1 hpi, and the apical side was washed. Cultures were further incubated at the indicated temperature. At the indicated time, apical virus release was assessed by plaque titration (**a,b**). Data represent the mean ± 95% CI of hAEC cultures from three different human donors. Individual points represent the average of two technical replicates. Values at 0 hpi indicate the titer of the inoculum used to infect the hAEC cultures, and values at 1 hpi indicate the remaining titer after the third wash. The p-values were computed by using two-sided paired sample t tests. At 96 hpi, hAEC cultures pretreated with 100 IU/ml type I IFN or 100 ng/ml type III IFN were fixed and processed for immunofluorescence analysis using antibodies against SARS-CoV Nucleocapsid protein (green), β-tubulin (cilia, red), ZO-1 (tight junctions, white) and DAPI (blue) (**c**). Representative z-projections of one donor are shown. Scale bar, 20 microns.

The reduction of viral progeny titers in both IFN pretreatment conditions were corroborated by immunofluorescence analysis at 72 hpi. Viral antigens were no longer detected upon immunostaining of the type I or III IFN-pretreated hAEC cultures with anti-SARS-CoV Nucleocapsid protein antibodies (**Fig. 2c**). Altogether, these results suggest that the viral replication kinetics of both SARS-CoV and SARS-CoV-2 in the upper and lower airways are heavily dependent on innate immune responses elicited by type I and III IFN, and that a potent IFN response can efficiently restrict viral replication of SARS-CoV-2 in primary well-differentiated hAEC cultures.

### Host response to SARS-CoV and SARS-CoV-2 infection in hAEC cultures at 33°C and 37°C

The notable impact of incubation temperature on SARS-CoV and SARS-CoV-2 replication kinetics (**Fig. 1**) and reduction in viral loads following both type I and III IFN pre-treatment (**Fig. 2**), prompted the assessment of host transcriptional response dynamics to SARS-CoV and SARS-CoV-2 infections at 33°C and 37°C. Cellular RNA was extracted from hAEC cultures infected with either SARS-CoV or SARS-CoV-2 (MOI 0.1) and from uninfected hAEC cultures at 24, 48, 72, and 96 hpi. The latter was then used for transcriptomics analysis following the Bulk RNA Barcoding and sequencing (BRB-seq) protocol ^30^. Data from seven different biological donors were collected and used to perform pairwise comparisons between either SARS-CoV or SARS-CoV-2 virus-infected hAEC cultures and unexposed hAEC cultures using three different approaches. First, pairwise comparisons were performed by segregating sample by both temperature and time prior to analysis. This approach identified a total of 185 differentially expressed genes (DEGs) for SARS-CoV and 351 DEGs for SARS-CoV-2 (Log_2_FC ≥ 1.5, FDR ≤ 0.1), represented by a total of 401 unique genes (**Supp. Fig. 3a-d, Supp. Table 1**).

To increase statistical power and identify DEGs that may have very low levels of expression in uninfected hAEC cultures, two further approaches to differential expression analysis were also performed. The first, which grouped samples by temperature, and then compared SARS-CoV or SARS-CoV-2 virus-infected hAEC cultures to uninfected hAEC cultures, identified a total of 126 DEGs for SARS-CoV at 33°C, 2 DEGs for SARS-CoV at 37°C, 161 DEGs for SARS-CoV-2 at 33°C, and 82 DEGs for SARS-CoV-2 at 37°C (**Fig. 3a, Supp. Table 2**). Hierarchical clustering of these SARS-CoV and SARS-CoV-2 DEGs uncovered that irrespective of temperature and time, the overlap between the host response induced by each virus was relatively low and the majority of upregulated DEGs were present at 72 and 96 hpi (**Supp. Fig. 3a-d**). Interestingly, many DEGs identified at 72 and 96 hpi for SARS-CoV-2 showed increased expression levels as early as 48 hpi at 37°C, but not at 33°C (**Fig. 3b**). Comparison of DEGs identified at each temperature for SARS-CoV-2 revealed a core group of 45 common upregulated ISG-related genes, including *IFNL1, CXCL10, CXCL11, IRF7, STAT1, IFI35, OASL, CMP2K, HELZ2, MX1, DDX60, IFI44L, ISG15*, and *IFIT3* (**Supp. Table 3**). Finally, pairwise comparisons were also performed for ungrouped samples between SARS-CoV or SARS-CoV-2 virus-infected hAEC cultures and uninfected hAEC cultures (**Supp. Table 4**). In addition to the individual comparisons for SARS-CoV-2, we also performed direct pairwise comparisons between SARS-CoV and SARS-CoV-2. The latter revealed that the most contrasting differences observed between the two viruses occurred at 96 hpi for 33°C, whereas they occurred at 72 hpi for 37°C (**Supp. Fig. 3e, f**). DEGs identified in the SARS-CoV-2 versus SARS-CoV analysis using the aforementioned three approaches are summarized in **Supp. Tables 5-7**.

**Figure 3.**
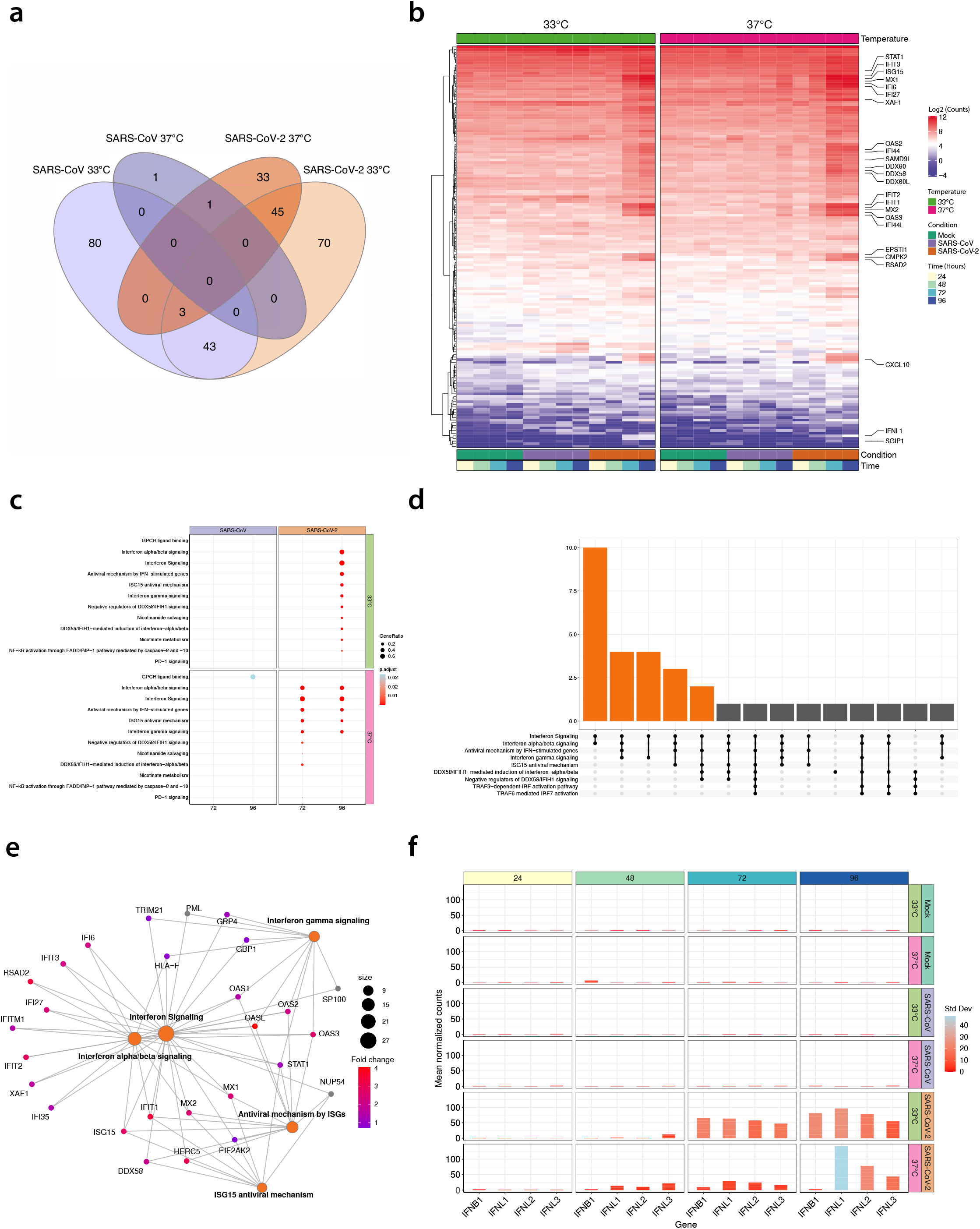
Temperature-dependent host transcriptional response in SARS-CoV and SARS-CoV-2 virus-infected hAEC cultures. Venn diagram showing the overlap among differentially expressed genes (DEG) identified in SARS-CoV (purple) and SARS-CoV-2 (orange) virus-infected hAEC cultures at either 33°C or 37°C (**a**). Hierarchical cluster analysis of DEGs identified in SARS-CoV and SARS-CoV-2 virus-infected hAEC cultures at either 33°C or 37°C compared to uninfected hAEC cultures (Mock). Expression levels for individual DEGs are shown in rows as the log_2_ mean normalized counts for seven human donors stratified by condition, temperature, and hours post-infection (columns; representative colours shown in legends). The top 25 DEGs for SARS-CoV-2 (ranked by adjusted p-values) are shown on the right (y-axis) (**b**). Dotplot illustrating pathway enrichment analysis performed on the 16 distinct DEG profiles. Significantly enriched pathways for SARS-CoV and SARS-CoV-2 are shown for both 33°C and 37°C incubation temperatures at 72- and 96-hours post-infection (hpi). Dots were adjusted in size and colour to illustrate the gene ratio and adjusted p-value (< 0.05) for a given pathway, respectively (**c**). An UpSet plot visualizing the intersection of the number of individual genes (y-axis) found to be associated with the significantly enriched pathways (x-axis). Intersections with 2 or more overlapping genes are highlighted in orange (**d**). Gene-concept network plot illustrating the individual relationships between DEGs and the top 5 significantly enriched pathways (orange) for SARS-CoV-2. Enriched pathway hubs (orange) were adjusted in size to reflect the number of genes associated with each respective pathway, whereas individual genes are coloured based on their expression level (fold change relative to expression level in uninfected hAEC cultures) (**e**). Bar graph illustrating the log2 mean normalized expression levels over time for *IFNL1, IFNL2, IFNL3*, and *IFNB1*, at the respective temperatures for Mock (uninfected) hAEC cultures (top two panels) and for SARS-CoV (middle two panels) and SARS-CoV-2 (bottom two panels) virus-infected hAEC cultures. Bars were adjusted in colour to illustrate the respective standard deviation (SD) among donors (**f**).

To investigate the hierarchical clustering results from the SARS-CoV or SARS-CoV-2 versus uninfected hAEC cultures in more detail, we annotated the top 25 upregulated DEGs for SARS-CoV-2 among the 16 distinct conditions (**Fig. 3b**). This revealed a clear temporal and temperature-dependent host response profile for SARS-CoV-2 virus-infected hAEC cultures, a finding that was less apparent for SARS-CoV. Furthermore, we found that *CXCL10, CXCL11*, and *TNFSF13B*, chemokines responsible for immune cell recruitment to the site of infection from the bloodstream, were among the core group of 45 common upregulated DEGs identified at both temperatures, albeit upregulation was stronger at 37°C compared to 33°C. Conversely, no upregulation was observed during SARS-CoV-2 infection in hAEC cultures for canonical pro-inflammatory genes such as *TNF, IL-11*, *IL-18, IL-6*, and *IL1B* (**Supp. Fig. 4a-b**). The pattern recognition receptor (PRR) *RIG-I/DDX58* and interferon-inducible 2’-5’-oligoadenylate synthetase-like protein (*OASL*), were also identified as some of the most prominent DEGs that showed an earlier and higher expression level at 37°C (**Fig. 3b**).

To establish whether any biological pathways were significantly enriched over time at the different ambient temperatures, pathway enrichment analysis was performed on all unique DEGs detected in the comparison analyses for SARS-CoV or SARS-CoV-2. Similar to the hierarchical clustering results, this analysis displayed a distinct temperature-dependent profile for SARS-CoV-2 at 72 and 96 hpi, specifically in diverse IFN and antiviral signalling pathways (**Fig. 3c**). Notably, none of these antiviral pathways were significantly enriched during SARS-CoV infection. The 43 unique genes associated with these enriched pathways, including *RIG-I/DDX58* and *OASL*, all displayed a clear temporal and temperature-dependent expression pattern that was inversely associated with the temperature-dependent viral kinetics difference for SARS-CoV-2 (**Fig. 3e, Fig. 1a, b, and Supp. Fig. 5**). Several individual genes were found to be associated with multiple enriched pathways, including canonical ISG-related genes such as *OASL, STAT1, ISG15*, and *IFIT1*. The intersection among the top 5 significantly enriched pathways for SARS-CoV-2 at 96 hpi for both temperatures, and 72 hpi for 37°C, is shown in **Fig. 3d**. A high degree of interconnectivity was observed among DEGs associated with these 5 pathways, further highlighting the importance of the host antiviral response during SARS-CoV-2 infection (**Fig. 3e**).

We previously observed, in the context of influenza A/H1N1 virus, that only a small fraction of infected cells produces IFN during infection ^31^. Since the majority of upregulated DEGs during SARS-CoV-2 infection were genes induced downstream of the IFN pathway, we next sought to assess the expression of individual type I and III IFN genes over time. Interestingly, although only *IFNL1* and *IFNB1* were classified as significantly upregulated DEGs in our analysis, the expression levels of *IFNL2* and *IFNL3* also followed a similar temperature-dependent pattern as the ISGs highlighted in **Fig. 3f**. In contrast, and in agreement with previous results (**Fig. 3a, e**), SARS-CoV infection did not induce type I or III IFNs at either 33°C or 37°C (**Fig. 3f**). Notably, we confirmed the expression of both type I and III IFN receptors (*IFNAR1, IFNAR2, IFNLR1*, and *IL10RB*) in multiple cell types of non-infected hAEC cultures using scRNA-seq (**Supp. Fig. 2e-h**). Together these results suggest that SARS-CoV infection triggers only very mild IFN induction in hAEC cultures, whereas SARS-CoV-2 infection leads to stronger induction of multiple IFNs that is dominated by type III IFNs and dependent on temperature ^31^.

Taken together, these results clearly demonstrate that SARS-CoV-2 and SARS-CoV induce disparate, virus-specific, and temperature-dependent host responses that inversely correlate with the previously observed dissimilar viral replication efficiencies at temperatures corresponding to the upper and lower respiratory tract. The majority of DEGs are related to the antiviral and pro-inflammatory response, which is much more pronounced in SARS-CoV-2 rather than SARS-CoV virus-infection, and the delayed induction of these DEGs at 33°C coincides with the increased replication of SARS-CoV-2 at temperatures corresponding to the upper respiratory epithelium.

## Discussion

In the current study, we demonstrate that the ambient temperatures reminiscent of the conditions in the upper and lower respiratory tract have a profound influence on both viral replication and virus-host dynamics, particularly innate immune responses, during SARS-CoV and SARS-CoV-2 infection in human airway epithelial cells. Using an authentic *in vitro* model for the human respiratory epithelium we demonstrated that SARS-CoV-2, in contrast to SARS-CoV, replicated up to 10-fold more efficiently at temperatures encountered in the upper respiratory tract. Concordantly, significantly increased amounts of Nucleocapsid-antigen positive cells were detected in these conditions. In addition, and despite intrinsic donor-to-donor variations, SARS-CoV-2 and SARS-CoV were highly sensitive to pretreatment with exogenous type I and type III IFNs. Importantly, temporal transcriptome analysis showed a temperature-dependent induction of the IFN-mediated antiviral and pro-inflammatory responses that inversely correlated with the observed replication kinetic efficiencies of both SARS-CoV and SARS-CoV-2 at temperatures found in the upper and lower respiratory tract.

One of the most profound phenotypical characteristics of fulminant SARS-CoV-2 is the early replication in the upper respiratory tract of infected individuals, which might facilitate the high transmissibility of SARS-CoV-2 ^14,24,25,32^. In contrast, SARS-CoV was shown to primarily replicate in the lower respiratory tract and efficient transmissibility occurred at later stages of the clinical course ^10,12,13^. Additionally, SARS-CoV was demonstrated to inhibit the induction of interferon responses in infected cells ^33,34^. The data presented here reflect these aspects of SARS-CoV and SARS-CoV-2 infections and contribute to the understanding of the disparate human-to-human transmission dynamics for both zoonotic coronaviruses. They provide a framework to address the parameters of the molecular basis of exacerbations induced by SARS-CoV-2 infection in predisposed individuals.

SARS-CoV-2 replication was strongly potentiated at 33°C and significant differences in the number of infected cells were observed between SARS-CoV and SARS-CoV-2 infected cultures. Given that the viral S protein receptor binding motifs interacting with the human receptor ACE2 are highly conserved between the two viruses and that both SARS-CoV and SARS-CoV-2 displayed a similar cell tropism, the 380 amino acid dissimilarities, distributed across the entire genome and distinguishing SARS-CoV-2 from SARS-CoV, might account for their differential replication efficiencies ^2,7–9^. Another factor that may have influenced our results is the 29-nucleotide truncation in the *ORF8* gene of SARS-CoV Frankfurt-1, which was maintained in the SARS-CoV lineage that initiated the international spread of SARS-CoV. Indeed, Muth and colleagues demonstrated that an intact *ORF8* invoked 10-fold higher replication kinetics at 37°C in various cell culture models, including hAEC cultures ^35^. Therefore, besides comparing the replication of different SARS-CoV *ORF8* variants at the temperature corresponding to the upper respiratory tract, it would be equally compelling to assess the phenotypic influence of similar truncations in the *ORF8* gene of SARS-CoV-2, especially since several SARS-CoV-2 isolates bearing a 382-nucleotide deletion truncating the *ORF8* gene have been detected ^36^. Such SARS-CoV-2 *ORF8* variants can be readily engineered using the reverse genetic systems that were recently established for SARS-CoV-2 ^32,37,38^.

In our study, we report different temperature-dependent viral replication efficiencies for SARS-CoV and SARS-CoV-2, inversely associated with the amplitude of the innate immune response, albeit with a more pronounced phenotype for SARS-CoV-2. Our results are consistent with other reports using undifferentiated primary cell-based systems where the induction of interferon responses was detected between 24 and 48 hours post infection with SARS-CoV-2 ^39^ In our study we analysed SARS-CoV-2-infected differentiated AEC cultures derived from multiple donors until 96 hpi at two different ambient temperatures. These data demonstrate a substantial induction of interferon and inflammatory responses in the airway epithelium following SARS-CoV-2 infection, in line with recent single-cell sequencing experiments performed on patient-derived samples ^40^. Foxman and colleagues elegantly described, in an analogous model to the human AEC cultures and by using common cold viruses, that the PRR-mediated IFN response is influenced by temperature ^23^. This may also apply in the context of SARS-CoV-2 infections, however, due to the multifaceted intricate nature of virus-host interactions it is likely that the efficient replication of SARS-CoV-2 and the concurrent expression of a plethora of known coronavirus antagonists of the antiviral response also play a crucial role herein ^41–46^. Nonetheless, we demonstrate that SARS-CoV-2 and SARS-CoV are sensitive to both type I and III IFN. These data are supported by several studies investigating the outcome of IFN pre-treatment in cultured cell lines ^39,47,48^ as well as the well-documented dominant antiviral role of type III IFN during virus infection in the respiratory epithelium ^31,49,50^ IFN lambda therefore represents an attractive candidate for the development of intervention strategies against SARS-CoV-2 respiratory infections.

The findings reported here provide crucial insight into the profound impact of ambient temperatures on pivotal virus – host interactions in the airway epithelium, which in turn influences the host immune response to SARS-CoV-2 infection ^51^. This work will likely lead to additional functional *in vivo* studies delineating the efficacy of antiviral responses triggered by SARS-CoV and SARS-CoV-2 infections, as well as deciphering the influence of viral antagonists and physical conditions. Therefore, the disparate viral replication efficiencies and host responses at 33°C and 37°C provide important insights into the molecular basis of SARS-CoV-2 infection and should be exploited broadly to support clinical interventions in COVID-19 patients.

## Material and methods

### Cells and human airway epithelial cell (hAEC) cultures

Vero-E6 cells (kindly provided by Doreen Muth, Marcel Müller, and Christian Drosten, Charité, Berlin, Germany) were propagated in Dulbecco’s Modified Eagle Medium-GlutaMAX supplemented with 1 mM sodium pyruvate, 10% (v/v) heat-inactivated fetal bovine serum (FBS), 100 μg/ml streptomycin, 100 IU/ml penicillin, 1% (w/v) non-essential amino acids and 15 mM HEPES (Gibco). Cells were maintained at 37°C in a humidified incubator with 5% CO_2_.

Primary human tracheobronchial epithelial cells were isolated from patients (>18 years old) undergoing bronchoscopy or pulmonary resection at the Cantonal Hospital in St. Gallen, Switzerland, or Inselspital in Bern, Switzerland, in accordance with ethical approval (EKSG 11/044, EKSG 11/103, KEK-BE 302/2015, and KEK-BE 1571/2019). Isolation and culturing of primary material was performed as previously described ^52^. Briefly, cryopreserved cells were thawed and expanded for one week in BEGM medium. After initial expansion phase, cells were transferred into in collagen type IV-coated porous inserts (6.5 mm radius insert, Costar) in 24-well plates. Cells were expanded for another 2-3 days in BEGM in a liquid-liquid state. Once the cells reached confluency, the basolateral medium was exchanged for ALI medium and the apical medium was removed to allow for the establishment of the air-liquid-interface (ALI). Basolateral ALI medium was exchanged three times per week and apical side was washed with Hanks balanced salt solution (HBSS, Gibco) once a week, until the development of a fully differentiated epithelium (3-4 weeks), which was monitored by optical microscopy. Several modifications to the original protocol were used. The concentrations of hydrocortisone for both BEGM and ALI were increased to 0.48 μg/ml and BEGM was further supplemented with the inhibitors 1 μmol/L A83-01 (Tocris, USA), 3 μmol/L isoproterenol (Abcam, Cambridge, United Kingdom) and 5 μmol/L Y27832 (Tocris, USA) ^53^. Basolateral ALI medium was exchanged three times per week and apical side was washed with hanks balanced salt solution (HBSS, Gibco) once a week. hAEC cultures were maintained at 37°C in a humidified incubator with 5% CO_2_.

### Viruses

SARS-CoV strain Frankfurt-1 (GenBank FJ429166) ^35,54^ and SARS-CoV-2 (SARS-CoV-2/München-1.1/2020/929) ^55^ were kindly provided by Daniela Niemeyer, Marcel Müller, and Christian Drosten, and propagated and titrated on Vero E6 cells.

### Infection of hAEC cultures

Well-differentiated hAEC cultures were infected with 30,000 plaque-forming units (PFU) of either SARS-CoV or SARS-CoV-2. Viruses were diluted in Hanks balanced salt solution (HBSS, Gibco), inoculated on the apical side and incubated for 1 h at 33°C or 37°C. Afterwards, virus inoculum was removed, and the apical surface washed three times with HBSS, whereby the third wash was collected as the 1 hpi timepoint. The cells were incubated at the indicated temperatures in a humidified incubator with 5% CO_2_. Released virus progeny were monitored every 24 h by incubating 100 μl of HBSS on the apical surface 10 min prior to the time point. The apical washes were collected, diluted 1:1 with virus transport medium (VTM), and stored at −80°C for later analysis. Basolateral medium was collected at each time point and stored at −80°C for later analysis. Fresh ALI medium was then added to the basolateral compartment. To analyze virus replication following interferon (IFN) exposure, hAEC cultures were pretreated with recombinant universal type I IFN (100 or 10 IU/ml; Sigma) or recombinant IFN-λ3 (100 or 10 ng/ml ^56^) for 18 h from the basolateral side, prior to infection and incubated at either 33°C or 37°C. As controls, untreated hAEC cultures were used. Shortly before infection with SARS-CoV and SARS-CoV-2, the basolateral medium containing type I or type III IFN was removed and replaced with medium without exogenous IFN.

### Immunofluorescence analysis of infected hAECs

Well-differentiated hAEC cultures were fixed with 4% (v/v) neutral buffered formalin and processed as previously described ^52^ Cells were permeabilized in PBS supplemented with 50 mM NH_4_Cl, 0.1% (w/v) Saponin and 2% (w/v) Bovine Serum Albumin (CB). To detect SARS-CoV and SARS-CoV-2, hAEC cultures were immunostained with a rabbit polyclonal antibody against SARS-CoV Nucleocapsid protein (Rockland, 200-401-A50), which also cross-react with SARS-CoV-2. Cell distribution of ACE2 were detected with a rabbit polyclonal antibody against ACE2 (ab15348, Abcam). Alexa Fluor^®^ 488-labeled donkey anti-rabbit IgG (H + L) (Jackson Immunoresearch) was used as secondary antibody. Alexa Fluor^®^ 647-labeled rabbit anti-β-tubulin (9F3, Cell Signaling Technology) and Alexa Fluor^®^ 594-labeled mouse anti-ZO1 (1A12, Thermo Fisher Scientific) were used to visualize cilia and tight junctions, respectively. Antibodies were diluted in CB. All samples were counterstained using 4’,6-diamidino-2-phenylindole (DAPI, Thermo Fisher Scientific) to visualize the nuclei. Samples were imaged on a DeltaVision Elite High-Resolution imaging system (GE Healthcare Life Sciences) equipped with 60x oil immersion objective (1.4 NA), by acquiring 200-300 nm z-stacks over the entire thickness of the sample. Images were deconvolved using the integrated softWoRx software. For the quantification of infected cells, images were alternatively acquired using an EVOS FL Auto 2 Imaging System equipped with a 20x air objective. All images were processed using FIJI software packages ^57^. Brightness and contrast were adjusted identically to their corresponding controls. Figures were assembled using the FigureJ plugin ^58^. Quantification of infected cells from four donors was performed by morphological segmentation of individual cells using the ZO-1 staining and the MorphoLibJ plugin in FIJI ^59^. Each region of interest was used to measure the mean intensity in the channel corresponding to the nucleocapsid staining. Cells with mean intensities > mean + 3 standard deviations compared to the distribution of mock-infected cells were considered positive. On average over 10^4^ cells were analysed per donor and per condition.

### Titration of apical and basolateral compartments

Viruses released in the apical or basolateral compartments were titrated by plaque assay on Vero-E6 cells. Shortly, 1.7*10^5^ cells/ml were seeded in 24-well plates one day prior to the titration and inoculated with 10–fold serial dilutions of virus solutions. Inoculums were removed 1.5 hpi and replaced with overlay medium consisting of DMEM supplemented with 1.2% Avicel (RC-581, FMC biopolymer), 5% heat-inactivated FBS, 50 mg/ml streptomycin and 50 IU/ml penicillin. Cells were incubated at 37 °C 5% CO_2_ for 48 hours, fixed with 4% (v/v) neutral buffered formalin and stained with crystal violet.

### Bulk RNA Barcoding and sequencing (BRB-seq) and data analysis

Total cellular RNA from mock and virus-infected hAEC cultures was extracted with the NucleoMag RNA kit (Macherey-Nagel) according manufacturers guidelines on a Kingfisher Flex Purification system (Thermofisher). Total RNA concentration was quantified with QuantiFluor^®^ RNA System (Promega) according manufactures guidelines on a Cytation 5 multimode reader (Biotek). A total of 100 ng of total cellular RNA was used for the generation of Bulk RNA Barcoding and sequencing (BRB-seq) libraries, and the subsequent sequencing on an Illumina HiSeq 4000 platform was done as previously described at a depth of approximately 12 million raw reads per sample ^30^. The sequencing reads were demultiplexed using the BRB-seqTools suite and were aligned against a concatenation of the human gene annotation of the human genome (hg38), SARS coronavirus Frankfurt 1 (AY291315) and SARS-CoV-2/Wuhan-Hu1/2020 (NC_045512) viral genomes using STAR and HTSeq for producing the count matrices. All downstream analyses were performed using R (version 3.6.1). ComBat-seq was used with default settings to adjust for batch effects in the raw data and generate an adjusted count matrix used for downstream analyses ^60^ Library normalization and expression differences between uninfected and virus-infected samples were then quantified using the DESeq2 package in R (version 1.28) with a fold change (FC) cut-off of ≥ 1.5 and a False Discovery Rate (FDR) of ≤ 0.1 ^61^. Due to the multi-factor design of these experiments, differential expression (DE) analysis was performed using several approaches: 1) Samples were subset by temperature and time prior to DE analysis (e.g. subset of samples at 3□C and 24 hpi). Infected samples were then compared to uninfected samples using the design ~ Batch + Condition to identify DEGs; 2) Samples were subset by temperature only prior to DE analysis (e.g. subset of samples for all time points at 3□C) and infected samples were compared to uninfected samples using the design ~ Batch + Condition; 3) Samples were kept together (not subset) and infected samples were compared to uninfected samples using the design ~ Batch + Condition. Substantial overlap was observed for DEGs identified using these 3 approaches and results for each approach are reported in Supplemental tables 1, 2, 4, 5, 6, 7. Venn diagrams were generated using the VennDiagram package in R for DEGs identified in approach 2 (**Fig. 3a**) and approach 1 (Supplementary Figure 3) ^62^.

Pathway enrichment analysis was performed using the clusterProfiler and ReactomePA packages in R ^63,64^. Significantly enriched pathways with a gene count > 1 and p-value of ≤ 0.05 were visualized using the enrichplot package. The complete list of significantly enriched pathways identified, as well as the genes associated with these pathways, are shown in Supplemental table 8. Additional data analysis and visualization was performed using a variety of packages in R, including ComplexHeatmap and ggplot2.^65^.

### Bioinformatic analysis of ACE2 and TMPRSS2 expression

For the analysis of *ACE2, TMPRSS2, IFNAR1, IFNAR2, IFNRL1*, and *IL10RB* mRNA expression we reanalysed the previously obtained single-cell raw sequencing data ^31^. The resulting unique molecule identifier (UMI) count matrix of each individual sample was pre-processed, filtered individually and merged in Seurat (v3.1). Data scaling, normalization and regressing out unwanted sources of variation (number of UMI’s, mitochondrial content, cell cycle phase) was performed using integrated SCtransform option, followed by dimensional reduction using UMAP (Uniform Manifold Approximation and Projection) embedding. For cell type annotation, the resulting integrated dataset was used for unsupervised graph-based clustering to annotate the different cell types using both cluster-specific marker genes and well-known canonical marker genes to match identified clusters with specific cell types found in the respiratory epithelium, as previously described^31^.

### Statistical testing

Distribution testing was performed using Shapiro-Wilk normality test (>0.05), followed by computing the P value of the mean log10 PFU/ml at each timepoint between SARS-CoV and SARS-CoV-2 using a two-sided paired sample t test. Analyses were performed using R (version 3.6.1).

### Data

Transcriptome data has been deposited in the Arrayexpress open-access public repository from the European Bioinformatics Institute (EMBL-EBI) under E-MTAB-9781, and scripts used for analysis and figure generation will be become available at Github upon publication.

## Acknowledgements

We gratefully thank the École Polytechnique Fédérale de Lausanne (EPFL) and the University of Bern for providing special authorization to conduct our research during the SARS-CoV-2 outbreak. We are grateful to Sabina Berezowska and Irene Ramos-Centeno (Institute of Pathology, University of Bern) for providing the tissues via the Tissue Bank Bern. This work was supported by the European Commission (Marie Sklodowska-Curie Innovative Training Network “HONOURS”; grant agreement No 721367), the Swiss National Science Foundation (SNSF) grants 179260, 160780, 173085, 31CA30_196644, the National Center of Competence in Research (NCCR) on RNA and Disease funded by the SNSF, and the German Federal Ministry of Education and Research, project RAPID.

## Author contributions

Conceptualization, R.D.; Investigation, P.V., M.G, S.S., J.K, J.R., B.M., E.C., J.P., M.H., A.K., L.L., M.W., J.P., T.T., N.E., H.S. and R.D., Resources, R.H., Writing – Original Draft, P.V. and R.D.; Writing – Review & Editing, P.V., J.K., V.T. and R.D.; Visualization, P.V., M.G., S.S., J.K. and R.D.; Supervision, V.G., D.A., B.D., V.T. and R.D.; Project Administration, R.D.; Funding Acquisition, V.T. and R.D.

