## Supplemental Figure Legends for "Disparate temperature-dependent virus – host dynamics for SARS-CoV-2 and SARS-CoV in the human respiratory epithelium"

**Supplemental figure 1. Basolateral release of SARS-CoV and SARS-CoV-2 in infected hAEC cultures.**

Well-differentiated hAEC cultures were infected with SARS-CoV and SARS-CoV-2 using 30,000 PFU or remain uninfected (mock), and were incubated at 37°C (**a**) or 33°C (**b**). Inoculated virus was removed at 1 hpi, and the apical side was washed. Cultures were further incubated at the indicated temperature. At the indicated time post infection, virus release in the basolateral compartment was assessed by plaque titration (**a-b**). Data represent the mean ± 95% CI of AEC cultures from three different human donors. Individual points represent the average of two replicates.

**Supplemental figure 2. ACE2 is expressed on both ciliated and non-ciliated cells**

Immunofluorescence analysis of ACE2 receptor distribution in unexposed well-differentiated hAEC cultures (**a**). Unexposed well-differentiated hAEC cultures were fixed and processed for microscopy analysis using antibodies against ACEC2 (green), β-tubulin (cilia, red), ZO-1 (tight junctions, white) and DAPI (blue). Representative z-projections of three donors are shown. Scale bar, 20 microns. Unexposed well-differentiated hAEC cultures from different human donors were used to perform scRNA-seq analysis. The UMPA plots shows the dimensional reduction of the 8128 cells belonging to either basal, goblet, secretory, preciliated and ciliated cell population (**b**), as well as the relative expression level and distribution of ACE2 (**c**) and TMPRSS2 (**d**), respectively. In addition, the respective relative expression level and distribution of both the interferon-alpha/beta receptor alpha and beta chain, IFNAR1 (**e**), IFNAR2 (**f**), and the type III IFN receptor complex IFNLR1 (**g**), and IL-10RB (**h**).

**Supplemental figure 3. Overlap of differential expressed genes in SARS-CoV and SARS-CoV versus SARS-CoV-2 virus-infected hAEC cultures**

Venn diagrams showing the overlap of differential expressed genes (DEG) in SARS-CoV or SARS-CoV-2 virus-infected hAEC cultures among the different time point and temperature conditions (**a-d**), and SARS-CoV-2 contrasted to SARS-CoV at the different time point and temperature conditions (**e-f**).

**Supplemental figure 4 Chemokine and Cytokine expression in untreated and virus-infected hAEC cultures.**

Bar graph illustrating the log2 mean normalized expression levels over time for the cytokines *IL11*, *IL18*, *IL1b*, and *TNF* (**a**), or the chemokines *CCL2*, *CCL5*, *CXCL10*, and *CXCL11* (**b**), at the respective temperatures for Mock (uninfected) hAEC cultures (top two panels) and for SARS-CoV (middle two panels) and SARS-CoV-2 (bottom two panels) virus-infected hAEC cultures. Bars were adjusted in colour to illustrate the respective standard deviation (SD) among donors

**Supplemental figure 5. Percentage viral reads in untreated and virus-infected hAEC cultures**

Boxplots illustrating the fraction of SARS-CoV (**a**) and SARS-CoV-2 (**b**) specific viral reads among the total fraction of sequence reads at the different time points and temperature conditions in the respective untreated (Mock), SARS-CoV and SARS-CoV-2 samples.
