## Supplementary figures and images for "Disparate temperature-dependent virus – host dynamics for SARS-CoV-2 and SARS-CoV in the human respiratory epithelium"

### Supplemental Figure 1

#### Supplemental figure 1

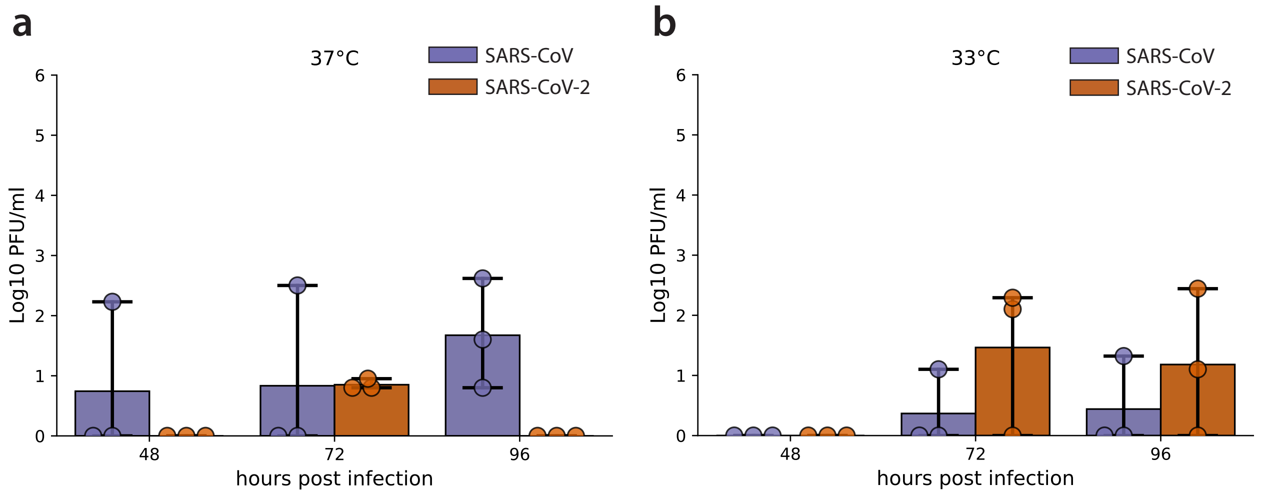

### Supplemental Figure 2

#### Supplemental figure 2

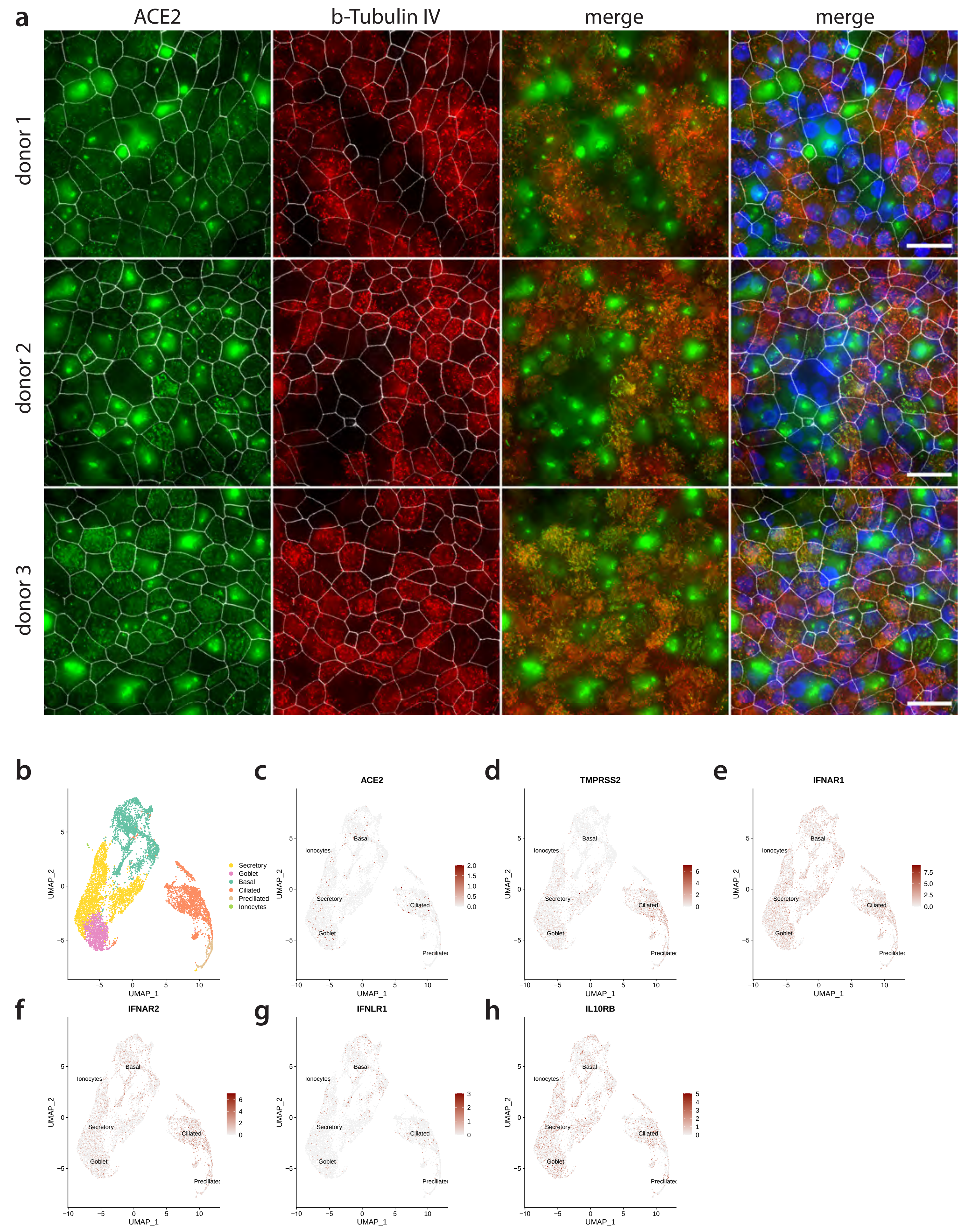

### Supplemental Figure 3

#### Supplemental figure 3

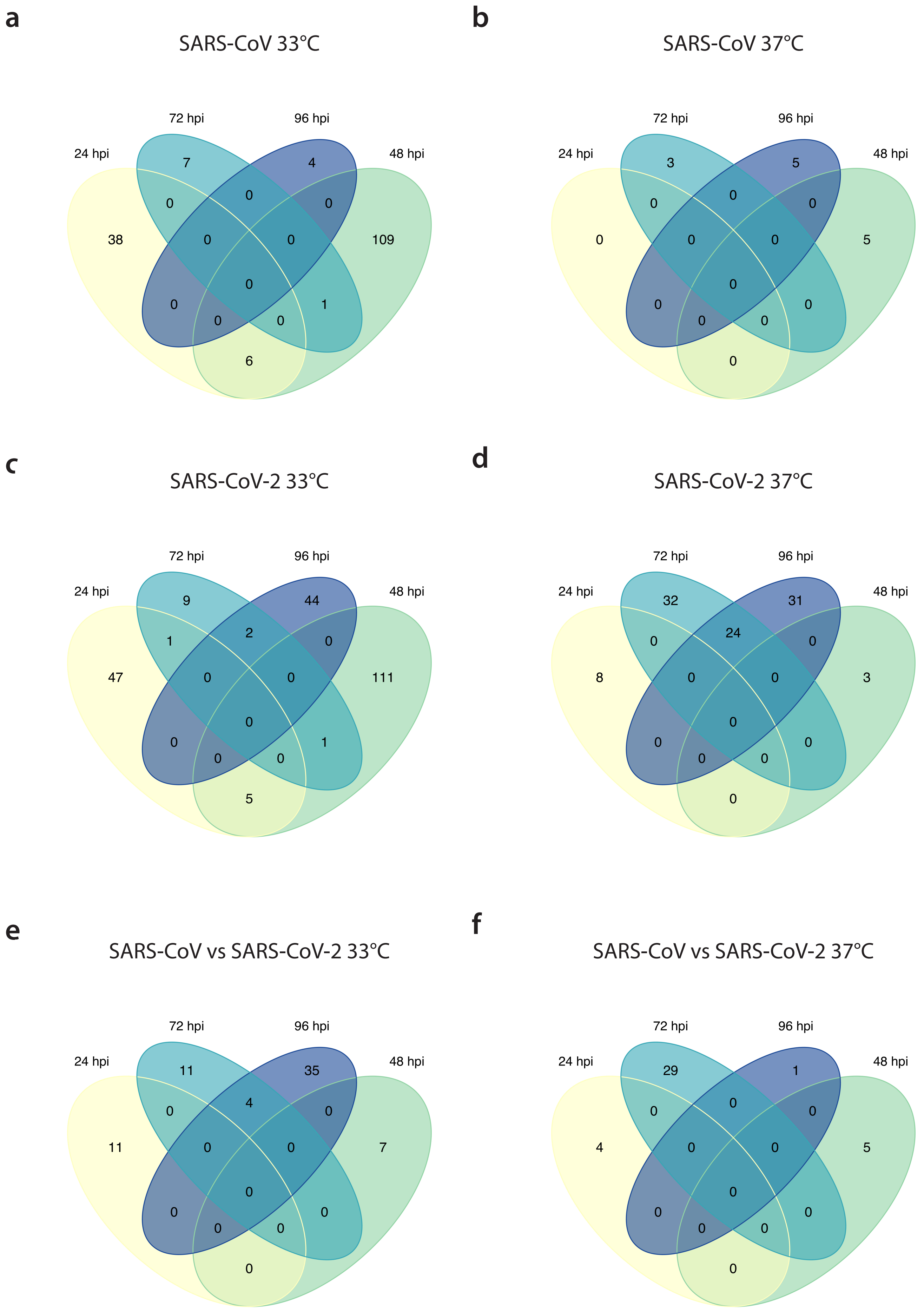

### Supplemental Figure 4

#### Supplemental figure 4

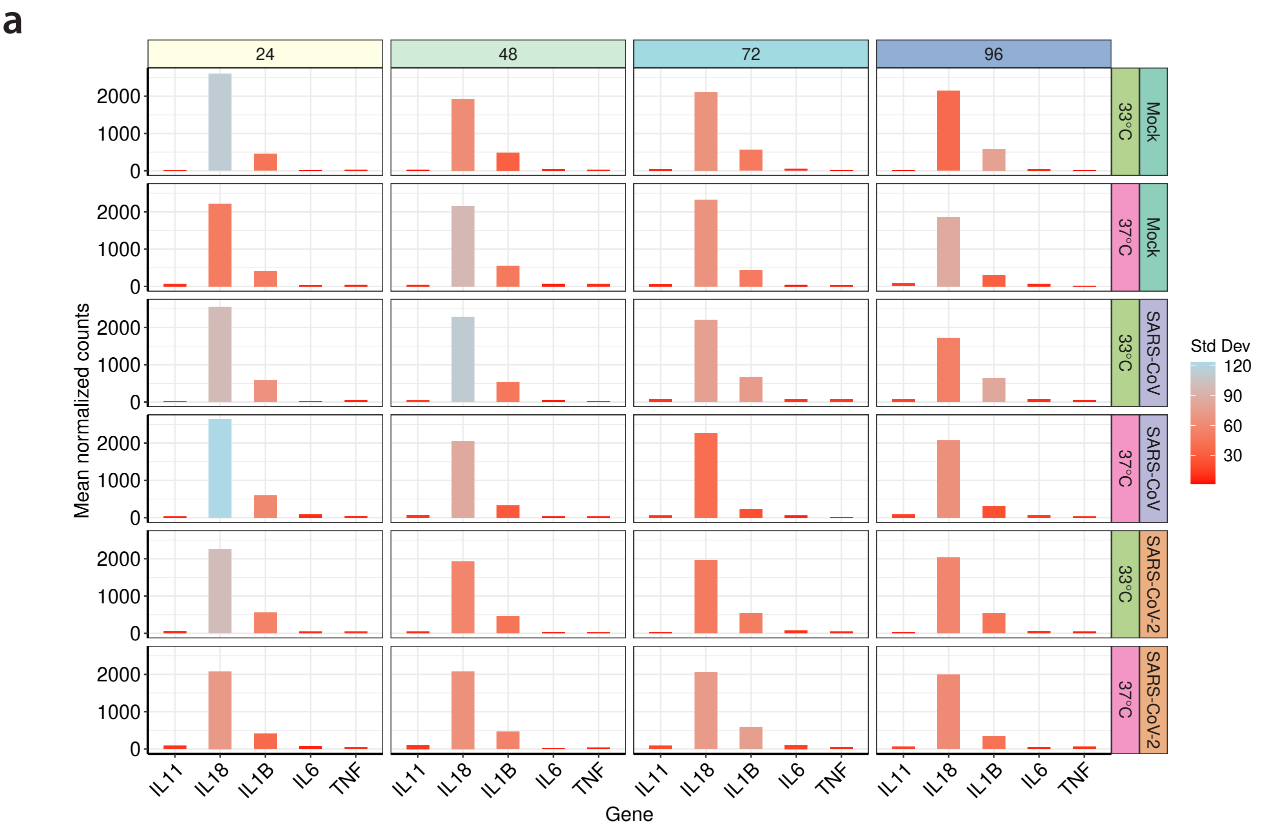


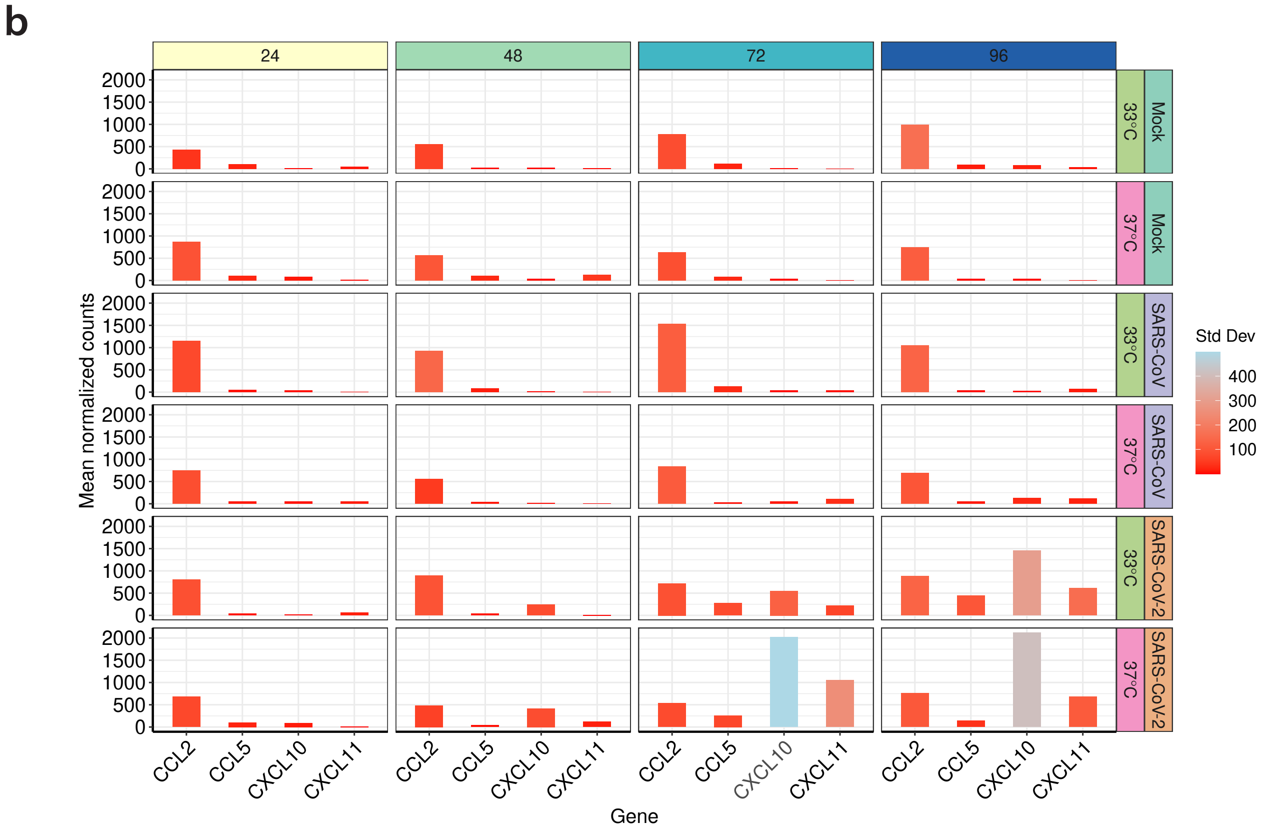

### Supplementary Figure 5

#### Supplemental figure 5

**a**

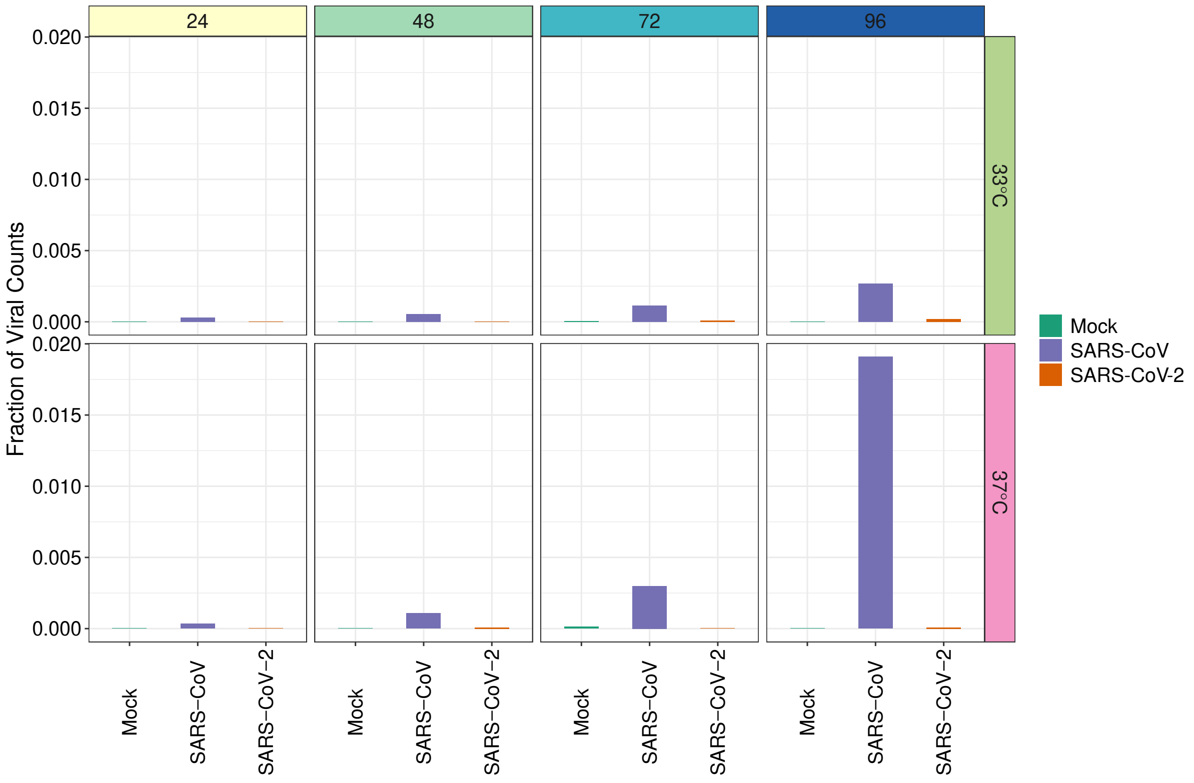


**b**

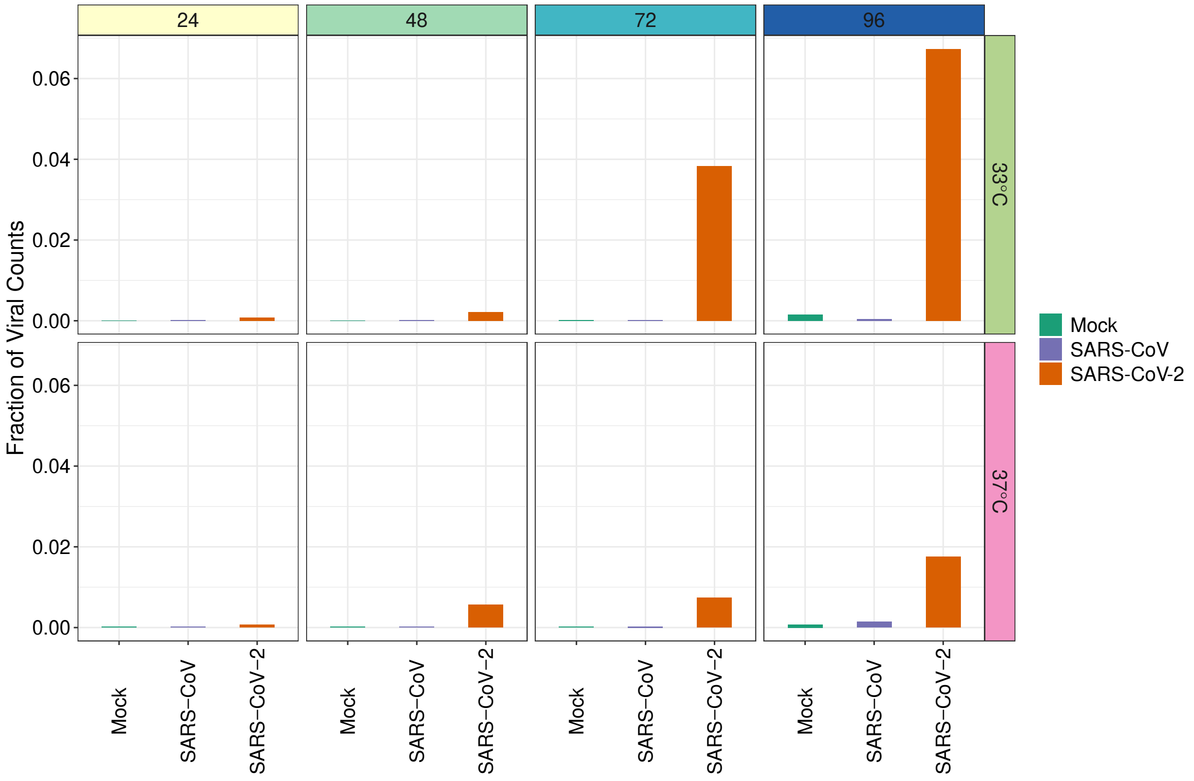
